# The context of experienced sensory discrepancies shapes multisensory integration and recalibration differently

**DOI:** 10.1101/2021.07.16.452674

**Authors:** Hame Park, Christoph Kayser

## Abstract

Whether two sensory cues interact during perceptual judgments depends on their immediate properties, but as suggested by Bayesian models, also on the observer’s a priori belief that these originate from a common source. While in many experiments this a priori belief is considered fixed, in real life it must adapt to the momentary context or environment. To understand the adaptive nature of human multisensory perception we investigated the context-sensitivity of spatial judgements in a ventriloquism paradigm. We exposed observers to audio-visual stimuli whose discrepancy either varied over a wider (± 46°) or a narrower range (± 26°) and hypothesized that exposure to a wider range of discrepancies would facilitate multisensory binding by increasing participants a priori belief about a common source for a given discrepancy. Our data support this hypothesis by revealing an enhanced integration (ventriloquism) bias in the wider context, which was echoed in Bayesian causal inference models fit to participants’ data, which assigned a stronger a priori integration tendency during the wider context. Interestingly, the immediate ventriloquism aftereffect, a multisensory response bias obtained following a multisensory test trial, was not affected by the contextual manipulation, although participant’s confidence in their spatial judgments differed between contexts for both integration and recalibration trials. These results highlight the context-sensitivity of multisensory binding and suggest that the immediate ventriloquism aftereffect is not a purely sensory-level consequence of the multisensory integration process.

## Introduction

Attributing a common source to two sensory signals, such as a sight and a sound, often results in their combination for perceptual judgements by the process of multisensory causal inference (Badde et al., 2020; Körding et al., 2007; Mihalik & Noppeney, 2020; Noppeney, 2021; Odegaard et al., 2015; Wallace et al., 2004). For example, when visual and acoustic cues are presented simultaneously from distinct locations the visual signal tends to influence judgements about the sound’s location when both seem to originate from a common source, such as during a ventriloquism act (Alais & Burr, 2004; Rohe & Noppeney, 2015b; Vroomen et al., 2001). However, when the spatial discrepancy between the two signals is large the multisensory judgement bias diminishes, as two very distant cues are unlikely to originate from a common cause.

As shown by Bayesian models, the interaction of two multisensory inputs can be decomposed into two contributions: one that depends on the physical discrepancy between the cues, and an overall binding tendency that is independent of the current stimulus (Beierholm et al., 2020; Körding et al., 2007; Odegaard et al., 2015; Park et al., 2021; Wozny et al., 2010). In typical laboratory studies, this overall binding tendency is often considered to be fixed for a given observer, reflecting their belief that the presented multisensory cues are causally related given the context of a specific experimental setting. Indeed, for a given task context the a priori binding tendency seems rather stable across days for individual observers (Odegaard & Shams, 2016). In everyday life, however, perception must adapt to momentarily encountered multisensory sensory discrepancies, in order to facilitate the context-sensitive binding of those cues that are indeed causally related. For example, when watching a movie on a TV with built-in speakers the spatial mismatch between the actor’s lips and her voice will be different than when viewing the same film using a surround-sound system or headphones. In real life, a given degree of spatial separation between two seemingly related signals may be surprisingly large in one context but not in another. Hence, two signals with the same degree of spatial separation may interact strongly during perceptual judgements in one context but not another. We here investigate this context-dependency of spatial multisensory perception using two prototypical multisensory judgement biases: the integration of two cues presented in the same trial and the immediate (trial-by-trial) aftereffect.

To study the context sensitivity of multisensory judgements we focused on an audio-visual spatial ventriloquism paradigm, which has proven to be a valuable window onto the computational and neural underpinnings of multisensory perception (Park & Kayser, 2019, 2021; Rohe & Noppeney, 2015a, 2016). Across two sessions, we presented audio-visual stimuli in two spatial contexts: one in which these stimuli covered a wide range of spatial discrepancies (between ± 46°) and one in which they covered only a narrower range (between ± 26°), while keeping the number of potential stimulus locations the same across contexts. By probing perceptual judgements about the stimulus location in both multisensory and unisensory trials we investigated two multisensory biases: that arising from the immediate combination of two simultaneously presented cues (the so-called integration or ventriloquism bias), and the aftereffect, which is visible in unisensory test trials following a previous multisensory stimulus (the so-called immediate ventriloquism aftereffect or recalibration bias (Bruns & Röder, 2015; Recanzone, 1998; Wozny & Shams, 2011b).

We expected that exposure to a wider range of spatial discrepancies would lead to an overall stronger integration bias. Specifically, in a context spanning a larger range of discrepancies a given discrepancy would be perceived as less extreme than in a context spanning only a smaller range, and this prominence of co-occurrence should foster the general belief in a common source for the visual and acoustic signals (Altieri et al., 2015; Ernst, 2007; Odegaard et al., 2017; Parise et al., 2012). This stronger belief about a common source then directly translates into a stronger integration bias for a given discrepancy, as predicted by Bayesian models of multisensory causal inference.

For the immediate aftereffect, we had no specific expectation as to how this would be affected by the contextual manipulation. Rather, we expected the data to arbitrate between two competing hypotheses. One line of work promotes the view that multisensory recalibration is directly linked to the causal inference process that moulds multisensory integration (Bruns & Röder, 2015; Noppeney, 2021; Park & Kayser, 2019). If this view is correct, the recalibration bias should adapt with context in a similar manner as the integration bias, hence resulting in a stronger bias for a wider contextual range of discrepancies in the present study. An alternative view stipulates that recalibration serves to increase perceptual accuracy by adapting unisensory representations contingent on the belief that these are subject to modality-specific biases, independent of whether two previous cues were deemed causally related or not (Di Luca et al., 2009; Noppeney, 2021; Zaidel et al., 2013). If this view is correct, one would not expect a difference in immediate recalibration bias between contexts.

## METHODS

### Participants

31 healthy right-handed adults (10 males, mean age 25.5 years, range 19 - 37 years) participated in this study. Following previous studies using similar experimental protocols (Park et al., 2021; Park & Kayser, 2020, 2021) and recommendations for sample sizes in empirical psychology (Simmons et al., 2011) we aimed for a sample size of at least n=20. Of the 31 recruited participants, three had to be excluded as they did not follow the task instructions (localizing the visual rather than the auditory target in AV trials) and two exhibited a below-threshold spatial localization performance during a screening task (see below). Data from 5 further participants had to excluded since they did not return for the second session. Therefore, here we report data from 21 participants (6 males, mean age 26.0 years, range 19 – 37 years). This study was approved by the local ethics committee of Bielefeld University. All participants submitted informed written consent and had self-reported normal vision and hearing and indicated no history of neurological diseases.

### Stimuli

The acoustic stimulus was a 1300 Hz sine wave tone (50 ms duration) sampled at 48 kHz and was presented at 64 dB r.m.s. through one of 5 speakers (Monacor MKS-26/SW, MONACOR International GmbH & Co. KG, Bremen, Germany), which were located at 5 horizontal locations. These locations differed between experimental contexts, which were manipulated on a session-by-session basis. In one context (‘WIDE’) the stimulus positions were: −23.2°, −11.6°, 0°, 11.6°, 23.2°, while in another context (‘NARROW’) these were −13.4°, −6.7°, 0°, 6.7°, 13.4°, vertical midline = 0°; Figure 1A). Sound presentation was controlled via a multi-channel soundcard (Creative Sound Blaster Z) and amplified via an audio amplifier (t.amp E4-130, Thomann Germany). Visual stimuli were projected (Acer Predator Z650, Acer Inc., New Taipei City, Taiwan) onto an acoustically transparent screen placed in front of the speakers (Screen International Modigliani, 2×1 m^2^). The visual stimulus was a cloud of white dots distributed according to a two-dimensional Gaussian distribution (200 dots, SD of vertical and horizontal spread = 1.6°, width of a single dot = 0.12°, duration = 50 ms), which has been used in previous studies on the ventriloquism effect and aftereffect (Park et al., 2021; Park & Kayser, 2020, 2021). The visual stimuli were centred around the same five locations as the acoustic stimuli (Figure 1B). Stimulus presentation was controlled using the Psychophysics toolbox (Brainard, 1997) for MATLAB (The MathWorks Inc., Natick, MA) which ensured temporal synchronization of auditory and visual stimuli.

**Figure 1.**
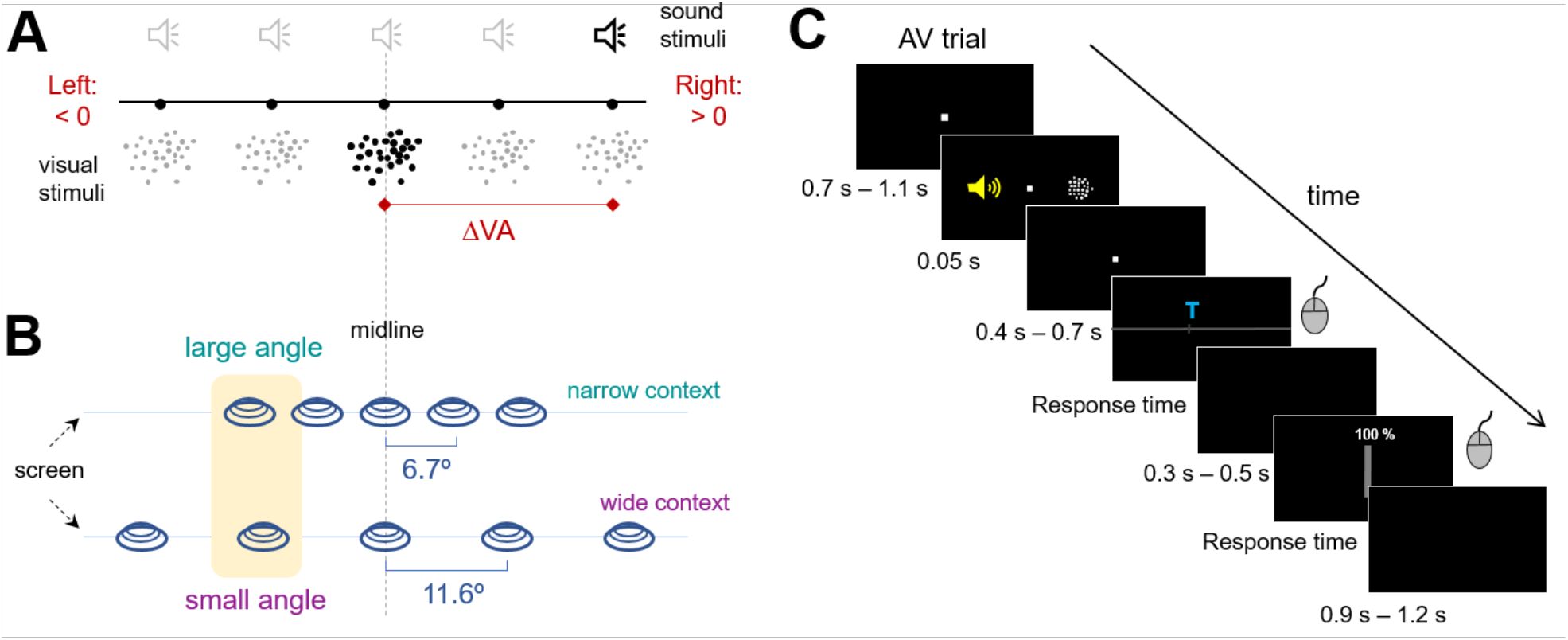
Stimuli and experimental task. (A) Auditory and visual stimuli were presented from five potential locations and were characterized by the respective unisensory locations and their spatial discrepancy (ΔVA). Sounds were presented from speakers occluded by an acoustically-transparent screen, on which the visual stimuli (clouds of dots) were projected. (B) In two separate sessions, these stimuli were presented in two contexts (NARROW, WIDE), characterized by a spacing of stimulus locations of either 6.7 ° (NARROW) or 11.6° (WIDE). Shaded area indicates how a similar angle can be interpreted as either large or small depending on the context. (C) The experiment consisted of audio-visual (AV) trials, auditory trials (A) and visual trials (V). Each trial had the same structure: after a random fixation period the stimulus was presented for 50 ms. After a post-stimulus period the participant was cued to indicate the perceived stimulus location by adjusting the position of a mouse cursor along the horizontal plane according to the perceived stimulus location. A visual cue (T: tone or V: visual) indicated which sensory modality had to be localized. After a second delay period participants were asked to indicate the confidence in their previous location rating using a vertical response bar.

### Experimental Setup

The paradigm was based on a single-trial audio-visual localization task (Park & Kayser, 2019; Wozny & Shams, 2011b), with trials and conditions designed to probe both the ventriloquism bias (integration) and the aftereffect bias (recalibration). Participants were seated in front of an acoustically transparent screen with their heads on a chin rest. The participants’ task was to localize a sound during either Audio-Visual (AV) or Auditory (A), trials, or to localize a visual stimulus during Visual (V) trials. These visual trials were included to keep the attention focused to both sensory modalities and to judge whether participants tended to localize the visual stimulus during AV trials (hence not following task instructions). We used an auditory report in the multisensory trials in line with previous studies (Bruns & Röder, 2015; Park & Kayser, 2019), and to reduce the length of the experiment in comparison to a dual report paradigm (Wozny & Shams, 2011b). During AV trials, the locations of auditory and visual stimuli were drawn semi-independently from the 5 locations to yield 9 different audio-visual discrepancies (abbreviated ΔVA). The range of physical discrepancies varied between contexts from either −46.4° to 46.4° (WIDE) or from −26.8° to 26.8° (NARROW), in 9 levels each. Each participant performed the experiment in the WIDE and NARROW contexts on a different day, and the order of the contexts were randomized across participants (half wide first, the other half narrow first). For each context, we repeated each audio-visual discrepancy 44 times. Each AV trial was followed by an A trial and 72 V trials were interleaved and always followed an A trial to not interrupt the AV-A sequence. For each context session, the experiment included 864 trials in total, which were divided into 4 blocks. Each trial started with a fixation period (uniform 700 ms – 1100 ms), followed by the stimulus (50 ms). After a random post-stimulus period (uniform 400 ms - 700 ms) the response cue emerged, which was a horizontal bar along which participants could move a cursor. A letter instructed participants about which stimulus they were supposed to localize (sound or visual stimulus). After a second delay (uniform 300 ms - 500 ms) we also obtained confidence ratings: a vertical bar with a cursor appeared, allowing participants to indicate their confidence on a scale of 0% - 100%. There was no constraint on response times. Inter-trial intervals varied randomly (uniform 900 ms - 1200 ms). A typical sequence of an AV trial is depicted in Figure 1C. Participants were asked to maintain fixation during the entire trial except the response, during which they could freely move their eyes.

Prior to the actual experiment each participant was tested for decent spatial hearing using a screening task. Here, participants were asked to localize noise bursts presented from the 4 lateralized speakers in a left/right two-choice task, performing 10 trials per location. Participants performing below 75% correct responses on average were excluded. Most participants were able to localize sounds well (average percent correct responses around 94% for either context).

### Model-free analyses of behavioural biases

Following previous work, we defined response biases as follows (Rohe & Noppeney, 2015b; Wozny & Shams, 2011a): the ventriloquism (integration) bias in AV trials was defined as the bias induced by the visual stimulus away from the true sound location: i.e. the difference between the reported sound location (R_AV_) and the location at which the sound (A_AV_) was actually presented (ve = R_AV_ - A_AV_). The aftereffect bias in the A trials was defined as the bias in the reported sound location relative to the audio-visual discrepancy experienced in the previous AV trial. This was computed as the difference between the reported sound location (R_A_) and the mean reported location for all A trials of the same stimulus position (μR_A_), i.e., vae = R_A_ - μR_A_. This was done to ensure that any overall bias in sound localization would not influence this measure, such as a tendency to perceive sounds closer to the midline than they actually are.

We used linear mixed models to quantify the dependencies of each bias on the audio-visual discrepancy (ΔVA) and to test for a potential effect of context (Park et al., 2021; Park & Kayser, 2020). Previous work has established that both the ve and vae bias can depend on the discrepancy ΔVA in a nonlinear manner. We hence first confirmed that this was the case for the present data as well. A linear dependency is predicted by models of multisensory fusion (Alais & Burr, 2004; Ernst & Bülthoff, 2004), while models of multisensory causal inference predict an additional a decay of the bias with increasing disparity (Cao et al., 2019; Körding et al., 2007; Rohe & Noppeney, 2015b). For each bias we determined its dependency on ΔVA by contrasting the predictive power of three candidate models, which were fit across all trial-wise biases of all participants and both contexts (using a Gaussian distribution and an identity link function):

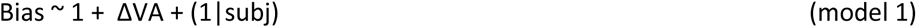

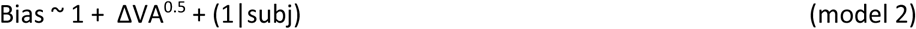

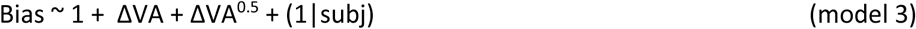

Here Bias is either ve or vae and subj indexes the participant as random effects. The square-root term (ΔVA^0.5^) describes the signed square-root of ΔVA (i.e. sign(ΔVA) * sqrt(abs(ΔVA))), and was chosen based on previous work (Cao et al., 2019; Park et al., 2021; Park & Kayser, 2020). Based on a group-level Bayesian Information Criterion (BIC) we found that the ve bias was best explained by model 3 (relative group-level BIC: 394, 9, 0 for models 1–3), while for the vae bias models 2 and 3 were equivalent (rel group-level BIC: 40, 0, 0). For reasons of parsimony, we relied on model 3 for both biases for subsequent analysis.

We then asked whether each bias was shaped by context, by adding interactions of each ΔVA-term with a (categorical) context variable, Cxt:

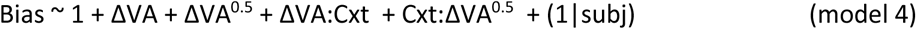

The parameter estimates for model 4 are provided in tables 1 and 3. There we also report Bayes factors for individual predictors, obtained by contrasting the full model with a reduced model omitting the respective predictor. In a control analysis we repeated the comparison of models 3 and 4 by focusing only on the levels of discrepancy that overlapped between wide and narrow contexts. For this, we excluded all trials with a discrepancy of ±46.4° or ±34.8° from the WIDE context data. Note that this resulted in unequal numbers of samples for each context (8316 for NARROW and 4620 for WIDE).

**Table 1.** *Generalized linear model analysis revealing a significant effect of context on the ventriloquism bias (ve). The model included a linear (ΔVA) and a nonlinear (ΔVA^0.5^) dependency on discrepancy and an interaction with context (Cxt, taking two categorical levels). Participants were included as random effects (subj). The table lists the parameter estimates (incl. confidence intervals), t- and p-values for the significance of each predictor and Bayes factors (BF) derived by contrasting the full model with reduced models omitting the specific predictor*. 16,632 observations from 21 participants.

| ve $\sim 1 + \Delta VA + \Delta VA^{0.5} + \Delta VA:Cxt + Cxt:\Delta VA^{0.5} + (1\vert subj)$ | | | | |
| --- | --- | --- | --- | --- |
|  | Estimate [CI] | t-value | p-value | BF |
| intercept | -0.64 [-1.47, 0.18] | -1.52 | 0.126 |  |
| $\Delta VA$ | -0.05 [-0.08, -0.02] | -3.42 | **0.001** | |
| $\Delta VA^{0.5}$ | 1.87 [1.70, 2.05] | 20.80 | **0** | |
| $\Delta VA:Cxt$ | 0.09 [0.03, 0.152] | 3.00 | **0.003** | -1.4 |
| $Cxt:\Delta VA^{0.5}$ | -0.78 [-1.07, -0.49] | -5.25 | **$2 \times 10^{-7}$** | $7.2 \times 10^3$ |

For the aftereffect we also probed whether adding the ve bias from the preceding trial to model 4 further improved model prediction (model 5), and whether this was further enhanced when adding an interaction of context and ve bias (model 6).

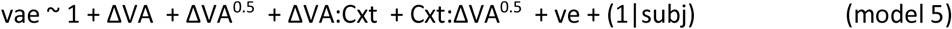

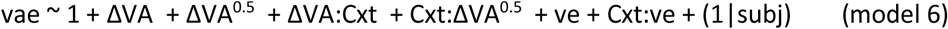

In a separate analysis we asked whether the distribution of the condition-wise and participant-wise (trial-averaged) biases show a significant pattern indicative of an effect of context (Park & Kayser, 2020). For this analysis we modelled the trial-averaged participant-wise biases against a combined linear and nonlinear dependency on ΔVA. We then used non-parametric tests to compare the linear and non-linear slopes separately between contexts.

### Analysis of confidence ratings

Confidence ratings delivered by the participants scaled between 0% and 100%. We then probed whether these confidence ratings depended on the eccentricity of the acoustic stimulus (coded as absolute deviation from the midline in degrees), on the absolute level of multisensory discrepancy (coded as the magnitude of the discrepancy) and on context. For this we again used linear mixed models with participants as random effects and context as categorical variable. For visualization in Figure 5 we standardized the condition-averaged (context, eccentricity, disparity) values within each participant separately for the AV and A trials, to remove between-participant fluctuations that would obscure the visualization of effects of interest.

### Model-based analysis of the ventriloquism effect

We used a Bayesian causal inference framework (Körding et al., 2007) to model the participant-wise ventriloquism biases obtained in the AV trials. Previous work has shown that this class of models captures the computations underlying flexible multisensory perception and can reproduce behavioural data well (Jones et al., 2019; Körding et al., 2007; Odegaard et al., 2015, 2017; Rohe et al., 2019; Rohe & Noppeney, 2015a; Wozny et al., 2010). These models reflect the inference about potential causal relations of two sensory signals based on prior experience and the available sensory evidence. To this end the model predicts the a posteriori probability of a single (C=1) or two distinct (C=2) causes underlying a pair of sensory signals by sampling from a binomial distribution with the common source prior p(C=1)=P_COM_ reflecting the overall probability of a common cause.

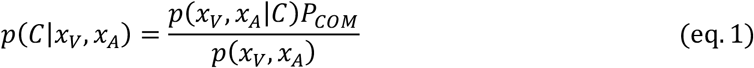

where *x_V_, x_A_* reflect noisy representations of visual and auditory inputs. In case of a common source underlying the two signals, the ‘true’ location (S) is drawn from a Gaussian spatial prior distribution with mean μ_P_ (here set to zero) and its standard deviation (σ_P_). In case of two independent sources, the true auditory and visual locations (S_A_, S_V_) are drawn independently from this prior distribution. These models incorporate sensory uncertainty by drawing the internal representations from independent Gaussian distributions centred on the true auditory (visual) locations, with given standard deviations (σ_A_, σ_V_). As detailed in several previous studies (Jones et al., 2019; Körding et al., 2007; Park et al., 2021; Wozny et al., 2010), the model then predicts optimal estimates of the auditory location for both the assumption of two separate causes or a common cause. In the latter case, the prediction is determined by the reliability-weighted average (Ernst & Banks, 2002), and the former case by the unisensory auditory representation combined with a central bias (the spatial prior). As in our previous work, we probed different decision functions to model the final estimate of the auditory location: Model Averaging (MA), Model Selection (MS), and Probability Matching (PM). We refer to previous work for details about these and the respective equations (Körding et al., 2007; Park et al., 2021; Rohe & Noppeney, 2015b). For a given decision function, the model comprises the following free parameters: the uncertainty in the sensory representations (σ_A_, σ_V_), the width of the spatial prior (σ_P_), and the a priori binding tendency (P_COM_).

To test the main question of whether multisensory causal inference adapts with the contextual manipulation, we fit two versions of these models. In one version we assumed that all model parameters were fixed for each observer across the two contextual manipulations (‘context-insensitive’ model). In a second version, we assumed that the sensory uncertainties and the width of the spatial prior remain the same across contexts, but allowed the a priori binding tendency to vary between contexts (‘context-sensitive’ model). We optimized the model parameters using the BADS toolbox based on the log-likelihood of the true data under the model (Bayesian Adaptive Direct Search (v1.0.3) (Acerbi & Ma, 2017). The lower and upper bounds for fitting the parameters were as follows: σ_A_ : 1°- 30°, σ_V_: 1°- 30°, σ_P_: 2°- 40° and P_COM_: 0.05 - 1. These values were chosen based on preliminary model fits that included an even larger range but revealed convergence within these boundaries for all participants.

The predicted conditional distributions for the spatial estimates for each model were obtained by marginalizing over the internal variables and were obtained by simulating the internal variables 20,000 times for each of the true stimulus conditions. To relate participants’ judgements to the model, both the actual data and the model predictions we binned both into 73 bins and we computed the log-likelihood of each participant’s actual data under a given model under the assumption that the different conditions are statistically independent. For each model and participant, we repeated the model fitting 500 times and selected the best run with the highest overall likelihood. We derived the participant-wise BIC values from the log-likelihoods and derived the group-level BIC value under the assumption that each participant adds independent evidence. We also quantified the model fit using the coefficient of determination R^2^ (Nagelkerke, 1991), calculated for each participant and model as 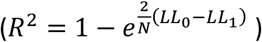. Here LL_1_ denotes the log-likelihood of the fitted model and LL_0_ that of a null model, defined as a model predicting a flat response distribution, and N is the number of data points used to fit the model. We verified that the model reflects the actual data well by simulating the predicted behavioural biases from the obtained model parameters for each participant based on 10,000 trials each (Palminteri et al., 2017).

In addition to modelling the participant-wise biases, we used these models to also extract an estimate of the predicted confidence. In the model, a Bayesian estimate of the confidence can be obtained as the probability that the predicted response is correct (Li & Ma, 2020). We here obtained this from the posterior probability distribution, which we evaluated at the value corresponding to the mean of the posterior, which we also used to generate the model prediction (Figure 3A). We used this definition, rather than e.g. the maximum of the posterior distribution, since for some decision functions the predicted distributions are not Gaussian. Predicted confidence ratings and localization biases were obtained for each participant using the best-fitting model parameters using the model selection decision function, as this decision function provided the best fit for the majority of participants. For visualization in Figure 5, we standardized the condition-averaged predicted confidence values within each participant similar to the actual data.

### Statistical Analysis

Confidence intervals (CI, 95%) were obtained using the bootstrap hybrid method with 199 resamples (Bootstrap Matlab Toolbox, Zoubir & Boashash, 1998). Linear mixed-effects models were fit using maximum-likelihood procedures (fitglme.m in Matlab), using a normal distribution and an identity link function. Bayes factors were computed as, where exp(BIC(H_0_) – BIC(H_1_)/2). We compared a model (H_1_) against the model minus a specific predictor of interest (H_0_), and report BF_10_ > 0 in favour of the model with the predictor (H_1_), and BF_10_ < 0 vice versa (Wagenmakers, 2007). The magnitudes of Bayes factors were interpreted using established conventions (Kass & Raftery, 1995). The dependency of confidence on discrepancy was analysed using a repeated measures ANOVA, as we had no clear expectation for a specific shape of the dependency of confidence on discrepancy. Bayes factors for the ANOVA were derived using the Bayes Factor Toolbox in Matlab (https://github.com/klabhub/bayesFactor). For non-parametric null hypothesis tests, we relied on sign-rank tests probing the null hypothesis of no difference in medians between contexts. For these we report p-values corrected for multiple comparisons across all sign-rank tests using the Benjamini & Yekutieli procedure (Benjamini & Yekutieli, 2001) and the respective test statistics, Z.

## RESULTS

Participants performed a modality-focused localization task of either auditory stimuli (presented in audio-visual or auditory trials) or visual stimuli (presented in visual trials). During each session these stimuli were presented from five independent locations (Figure 1A). Importantly, across two sessions presented on different days we manipulated the multisensory context by varying the degree of spatial discrepancies over either a narrow (± 26°) or a wide range (± 46°). This contextual manipulation was realized by manipulating the separation of the five potential stimulus locations, using a separation of 6.7° and 11.6° for the narrow and wide contexts respectively (Figure 1B). After each stimulus participants were cued as to which stimulus modality to localize and submitted their responses using a mouse cursor (Figure 1C). In addition, on each trial they were asked to rate their confidence for this localization judgement. For most parts of the experiments trials consisted of sequences of AV and A trials, which allowed us to probe the ventriloquism bias (ve) in the AV trial, which describes the mislocalization of the position of the sound resulting from the influence of the simultaneous visual cue. It also allowed us to probe the ventriloquism aftereffect (vae) during the A trials, which describes the mislocalization of a sound during a unisensory test trial following a multisensory stimulus.

### Effects of context on the integration bias

Figure 2A displays the group-level ventriloquism bias (ve) for each context as a function of the actual spatial disparity, and Figure 2B displays the data scaled to the same range for both contexts (n=21). As the figure suggests, the ve bias was significantly shaped by context. An analysis using linear mixed models showed that the ve bias depended both in a linear and a non-linear manner on multisensory discrepancy (ΔVA) (model 3; see Material and Methods). Based on this result, we then asked whether the group-level data was even better described by a model rendering the influence of discrepancy on bias context-dependent (model 4). This was indeed the case, and we found decisive evidence for an improved model fit when including interactions between context and the two ΔVA-terms as predictors (BF = 2×10^21^). As shown in Table 1, the interactions of both ΔVA-terms with context were significant.

**Figure 2.**
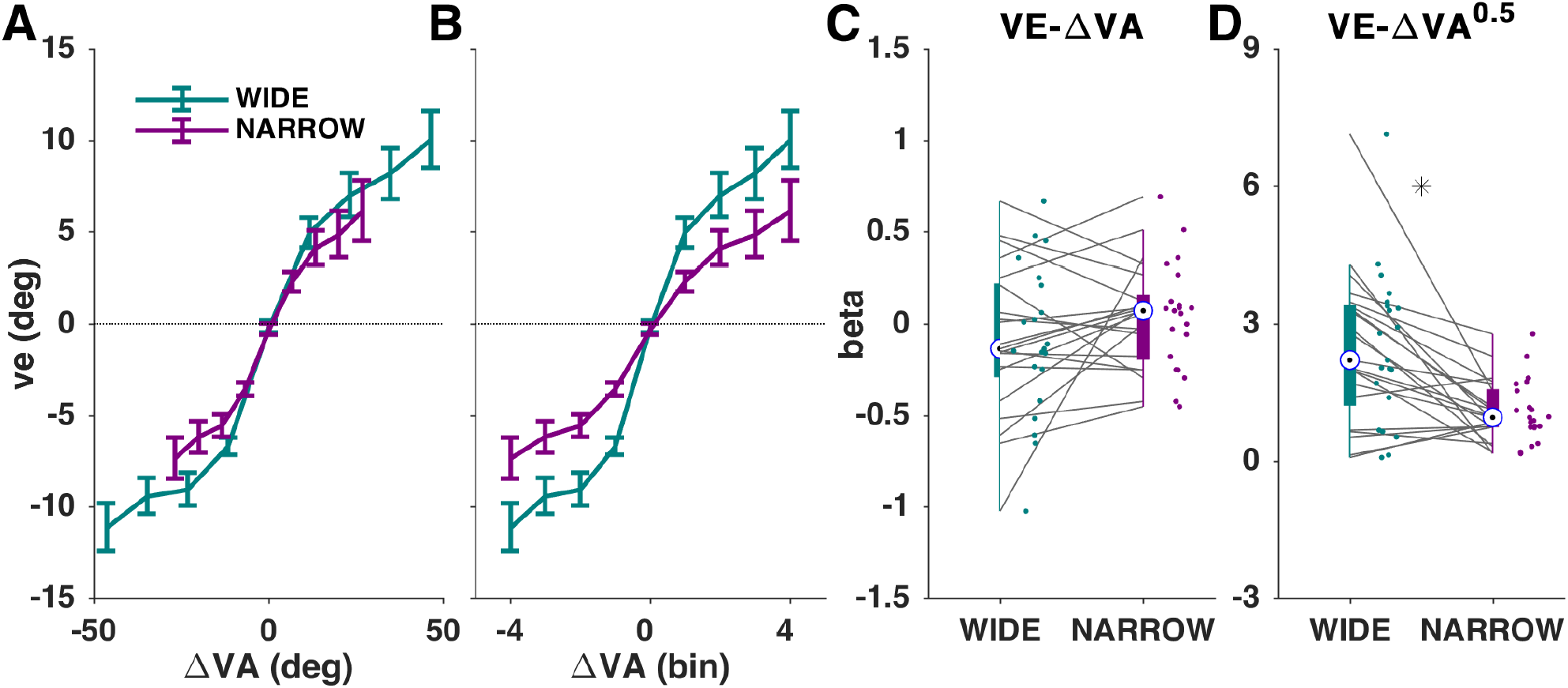
Context dependency of the integration bias in the AV trials. (A) The integration bias (ve) for each context as a function of the actual audio-visual discrepancy in degrees. The graph displays the group-mean and standard error of mean (n=21). (B) Same data shown when the ranges of discrepancies were normalized, i.e. showing the data as a function of the difference between the 5 discrete locations per experimental session. (C, D) Individual participant slopes describing the linear (C) and non-linear (D) dependency of ve on discrepancy (ΔVA). Asterisk denotes a significant difference (p < 0.05) based on a sign-rank test, corrected for multiple comparisons using the Benjamini & Yekutieli procedure. Teal: WIDE; Plum: NARROW.

To substantiate this result, we implemented a separate analysis in which we quantified the linear and non-linear dependencies of individual participant’s trial-averaged data against ΔVA (Figure 2C, D). We used non-parametric tests to probe for an effect of context (sign-rank tests, corrected for multiple comparisons for both ve and vae biases using the Benjamini & Yekutieli procedure (Benjamini & Yekutieli, 2001), which revealed a significant context effect for the nonlinear (Z=3.2, p_corr_= 0.0085) but not the linear slope (Z=−1.2, p_corr_=0.46).

Given that the two contexts covered distinct and partly non-overlapping ranges of discrepancies, we implemented a control analysis focusing only on the overlapping range of discrepancies. For this we excluded the two most extreme discrepancies (±46.4, ±34.8) from the WIDE context data. Still, a linear-model based analysis revealed decisive evidence that a model including context better predicted the data (BF = 3×10^26^) and the interaction of context and the nonlinear ΔVA-dependency was significant (β= −0.715 CI [−1.14, −0.28], t=−3.28, p =0.001, BF=1.9), while the interaction with a linear ΔVA-dependency was not (β = 0.07 CI [−0.03, 0.16], t= 1.35, p=0.16, BF=−42).

### Bayesian modelling of the ventriloquism bias

Previous work has shown that the ventriloquism bias is well captured by Bayesian causal inference models (Körding et al., 2007; Park et al., 2021; Rohe & Noppeney, 2015b). These describe the response bias by modelling the two unisensory representations using likelihood functions reflecting the respective sensory reliabilities (σ_A_, σ_V_). These unisensory representations are combined for a perceptual judgement based on an a priori binding tendency (P_COM_) and a suitable decision function (see Material and Methods and Figure 3). We here fit such models to individual participant data, comparing two models: one assuming the same binding tendency across contexts, and one allowing for a change in the a priori binding tendency between contexts, while keeping the other parameters fixed across contexts. Given that in this model the response bias results from the interaction of sensory uncertainties and an a priori binding tendency, we expected the context-sensitive model to explain the data better, and we expected the wider context to result in a higher binding tendency in the context-sensitive model. The model fitting results supported both hypotheses.

**Figure 3.**
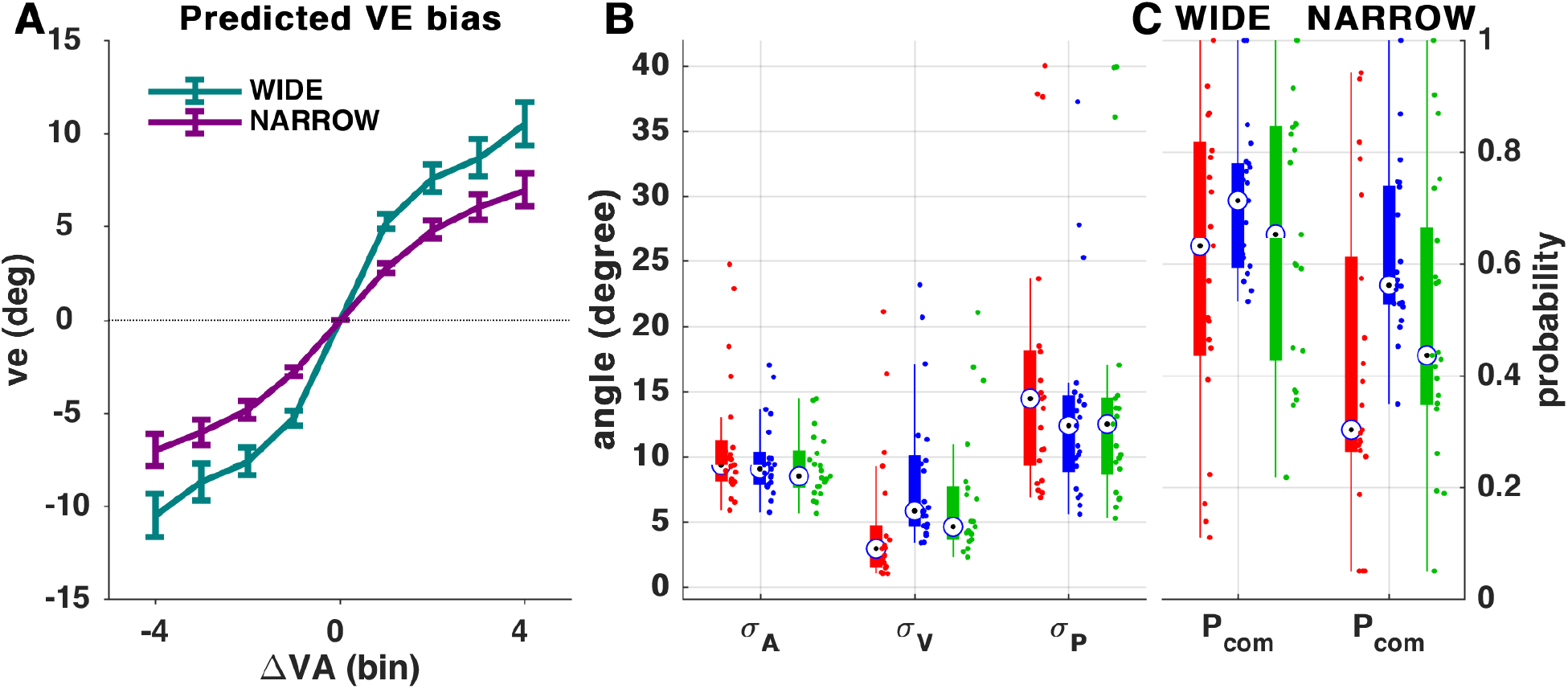
Bayesian causal inference models unveil a change in a priori binding tendency with context. (A) Predicted ve based on the parameters of the participant-wise best models (using MS as decision function). The bias is shown as a function of the difference in stimulus locations (similar as Figure 2B). (B,C) Model parameters for the context-sensitive model for each decision function shown as median (circle) and 25th to 75th quantiles (thick bars) and individual data (dots). Red: MA, Blue: MS, Green: PM.

Across participants we found that the models explained the individual-participant data well (see R^2^ values in Table 2; see Figure 3A for predicted biases). Importantly, a comparison of group-level relative BIC values provided decisive evidence that the context-sensitive model explained the data better (BIC difference context-sensitive minus context-insensitive models: 512, 248, 263, for the three decision functions of model averaging, MA, model selection, MS and probability matching, PM). Within the context-sensitive model the group-level BIC favoured model selection as best-fitting decision function (rel. BIC = 332, 0 and 21) while the model frequencies suggest that no decision function clearly prevailed across the group of participants (29%, 57% and 14%, respectively). Interestingly, the three decision functions implemented the experimentally observed ve biases by slightly different combinations of the visual reliability (σ_V_) and the a priori binding tendency (Figure 3B). For this reason, we contrasted the a priori binding tendency between contexts separately for each decision function. Non-parametric tests revealed significant or trending differences of P_COM_ between contexts (sign-rank tests, Z= 2.14, 1.93, 1.96 and p=0.032, 0.054, 0.049 for MA, MS and PM; Figure 3C), supporting that a change in multisensory spatial context leads to different a priori binding tendencies.

**Table 2.** Parameter estimates for the context-sensitive Bayesian causal inference models. Parameter estimates are provided for each decision function, shown as group-level median values and 95% bootstrap confidence intervals (brackets). The last column indicates the model fit (generalized R^2^). Decision functions: Model averaging (MA), Model selection (MS) and Probability matching (PM).

| | $\sigma_A$ | $\sigma_V$ | $\sigma_P$ | $P_{COM-wide}$ | $P_{COM-narrow}$ | $R^2$ |
| --- | --- | --- | --- | --- | --- | --- |
| MA | 9.35 [8.00, 10.4] | 2.91 [2.24, 4.29] | 14.6 [12.2, 19.2] | 0.63 [0.47, 0.80] | 0.30 [0.03 - 0.34] | 0.89 [0.85, 0.92] |
| MS | 9.06 [8.24, 10.1] | 5.82 [1.95, 6.85] | 12.4 [10., 15.5] | 0.71 [0.65, 0.81] | 0.56 [0.39 - 0.60] | 0.89, [0.86, 0.92] |
| PM | 8.49 [7.2, 9.28] | 4.60 [1.63, 5.16] | 12.5 [10.5, 15.9] | 0.65 [0.46, 0.85] | 0.44 [0.23, 0.52] | 0.89, [0.87, 0.91] |

### Effects of context on the aftereffect bias

We repeated the same analysis for the ventriloquism aftereffect bias (vae). Similar to the ve bias, the aftereffect increased with multisensory discrepancy, but without obvious differences between contexts (Figure 4A, B). When comparing models with or without context as a factor we found decisive evidence for no effect of context (BF=−665). The parameter estimates derived from the model including interactions between context and the discrepancy (model 4) revealed very strong evidence against any interaction of discrepancy and context (see p-values and Bayes factors in Table 3). In addition, the non-parametric analysis of individual-participant trial-averaged biases (Figure 4C, D) revealed no significant effect of context for either the linear (Z=−2.0, p_corr_=0.16) or the nonlinear slopes (Z=1.79, p_corr_=0.20). We again implemented a control analysis focusing on the range of discrepancies overlapping between the two contexts. Similar to the integration bias, this confirmed the results obtained using all levels of discrepancy: there was decisive evidence for no effect of context when comparing models (BF=−8×10^3^) and the parameter estimates for the model including context were not significant (interaction of context with the linear ΔVA-term: β = −0.01 CI [−0.07, 0.07], t=−0.11, p=0.90, BF=−113; the non-linear ΔVA-term: β = 0.05 CI [−0.28, 0.37], t=0.27, p=0.78, BF=−109).

**Figure 4.**
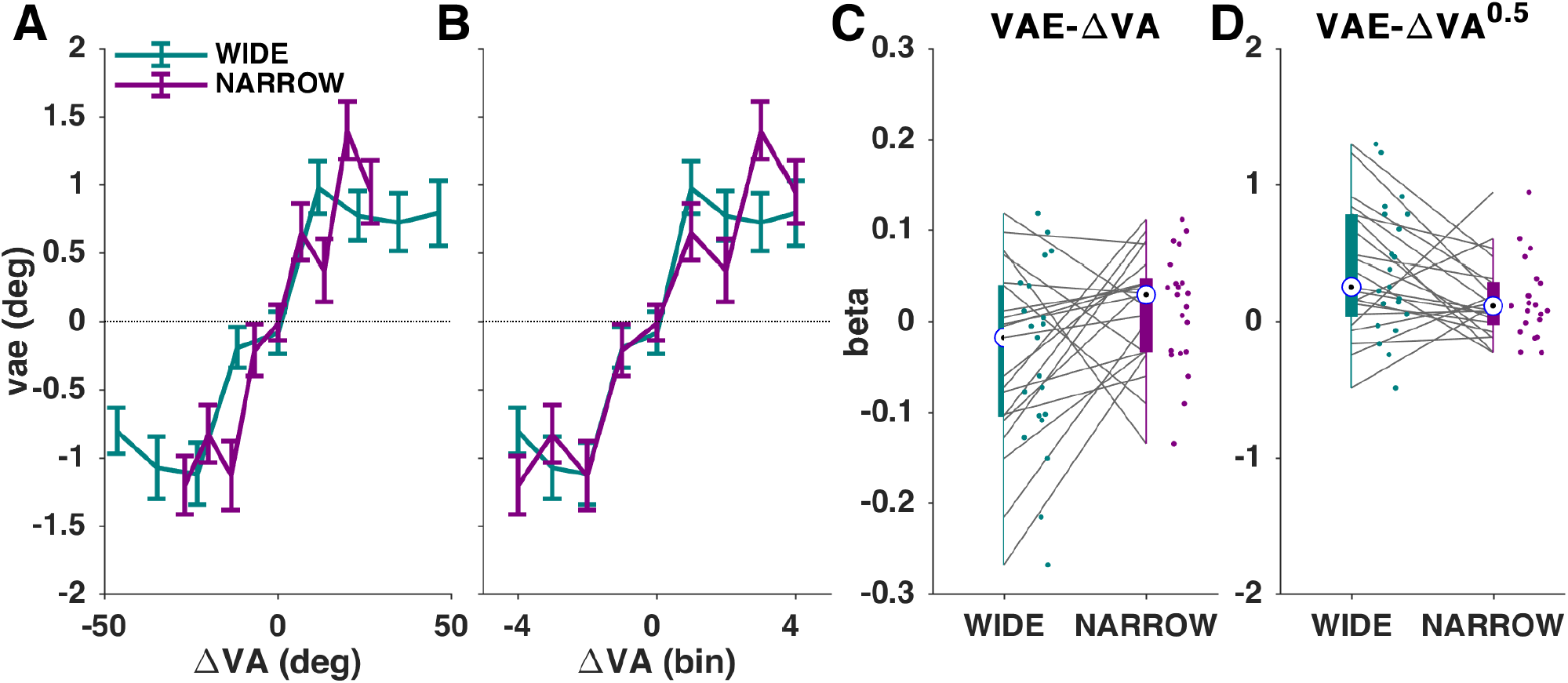
Context independency of the aftereffect bias obtained in the A trials. (A) The aftereffect bias (vae) for each context as a function of the actual audio-visual discrepancy in degrees. The graph displays the group-mean and standard error of mean (n=21). (B) Same data shown when normalized to the same range of discrepancies. (C, D) Individual participant slopes describing the linear (C) and non-linear (D) dependency of ve on discrepancy (ΔVA). Teal: WIDE; Plum: NARROW.

**Table 3.** *Generalized linear model analysis reveals no effect of context on the ventriloquism aftereffect (vae). The model included a linear (ΔVA) and a nonlinear (ΔVA^0.5^) dependency on discrepancy and an interaction with context (Cxt, taking two categorical levels). Participants were included as random effects (subj). The table lists the parameter estimates (incl. confidence intervals), t- and p-values for the significance of each predictor and Bayes factors (BF) derived by contrasting the full model with reduced models omitting the specific predictor*. 16,632 observations from 21 participants.

| vae ~ 1 + ΔVA + ΔVA<sup>0.5</sup> + ΔVA:Cxt + Cxt:ΔVA<sup>0.5</sup> + (1\|subj) |  |  |  |  |
| --- | --- | --- | --- | --- |
|  | Estimate [CI] | t-value | p-value | BF |
| intercept | -10<sup>-16</sup> [-0.08, 0.08] | -3x10<sup>-15</sup> | 1 |  |
| ΔVA | -0.02 [-0.04, -0.01] | -2.12 | **0.03** |  |
| ΔVA<sup>0.5</sup> | 0.28 [0.15, 0.41] | 4.36 | **2x10<sup>-5</sup>** |  |
| ΔVA:Cxt | 0.03 [-0.01, 0.08] | 1.63 | 0.10 | -34 |
| Cxt:ΔVA<sup>0.5</sup> | -0.12 [-0.33, 0.09] | -1.16 | 0.24 | -65 |

### Is the aftereffect directly related to the preceding integration bias?

One origin of the immediate aftereffect bias supposedly lies in the multisensory integration during the preceding audio-visual trial and the perceived common cause implicitly attributed to the auditory and visual cues (Badde et al., 2020; Noppeney, 2021; Rohlf et al., 2020). This belief in a common cause may carry over between trials and shape judgements of subsequent unisensory stimuli in proportion to the previous multisensory response bias, hence in the direction of the previous discrepancy. The data presented above support such a direct link between ve and vae biases, as both biases depend on the degree of audio-visual discrepancy in the AV trial. However, previous work has shown that the aftereffect can in part also be predicted by the response in the AV trial in addition to the sensory discrepancy (Park et al., 2021). In the following we characterize the relation between the two multisensory response biases and its dependency on the contextual manipulation in more detail.

First, we asked whether the aftereffect bias depends on the directly preceding ventriloquism bias, ve, in addition to the two predictors capturing the influence of the sensory discrepancy. Here, the ve bias serves as a direct indicator of the trial-wise integration of the multisensory information. Indeed, a comparison of models with and without the ve as an additional predictor revealed decisive evidence in favour of an influence of ve (model 4 vs. model 5; BF = 2×10^14^). We then asked whether ve as predictor interacts with context (model 6). This was not the case: while the main influence of ve on the aftereffect was highly significant (β= 0.05, CI [0.03, 0.06], t =6.5, p<10^−9^) the interaction with context was not significant and we obtained strong evidence for a null result (β: 0.004, CI [−0.018, 0.025], t=0.35, p=0.72, BF=−121). This corroborates previous work by demonstrating that the aftereffect is shaped by both the previous physical stimuli and the previous response. However, neither of these statistical links changes with context.

### Analysis of confidence data

We analysed subjective confidence ratings to understand whether this signature of meta-cognitive assessment is influenced by the contextual manipulation. As a sanity check, we first probed for an expected scaling of confidence with the difficulty of the auditory localization task, hence with the eccentricity of the auditory stimulus. The confidence ratings obtained in the AV trial directly confirmed this expected result, which was also predicted by the Bayesian causal inference model (Figure 5A, B): a linear model revealed a highly significant effect of eccentricity (β = 0.33, CI [0.28, 0.377], t=14.7, p ≅ 0). The addition of context as factor to the model revealed decisive evidence for an improvement in model fit (BF=223). While there was strong evidence against an effect of context itself (β = −0.38, CI [−1.4, 0.70], t=−0.67, p=0.50, BF=−102), we found anecdotal evidence in favour of an interaction (β = 0.15, CI [0.07, 0.24], t=3.44, p=0.001, BF=2.9), which reflected a stronger scaling of confidence per eccentricity in the narrow context. The confidence ratings in the A trial revealed same results: the inclusion of context as predictor significantly improved the model fit (BF=282), the effect of eccentricity was highly significant (β = 0.5, CI [0.47, 0.54], t=27.8, p ≅ 0) and we obtained substantial evidence for an interaction with context (β 0.13, CI [0.06, 0.20], t=3.6, p=0.0003, BF=5) and decisive evidence against an effect of context itself (β −0.23, CI [−1.0, 0.59], t=−0.55, p=0.57, BF=−110).

**Figure 5.**
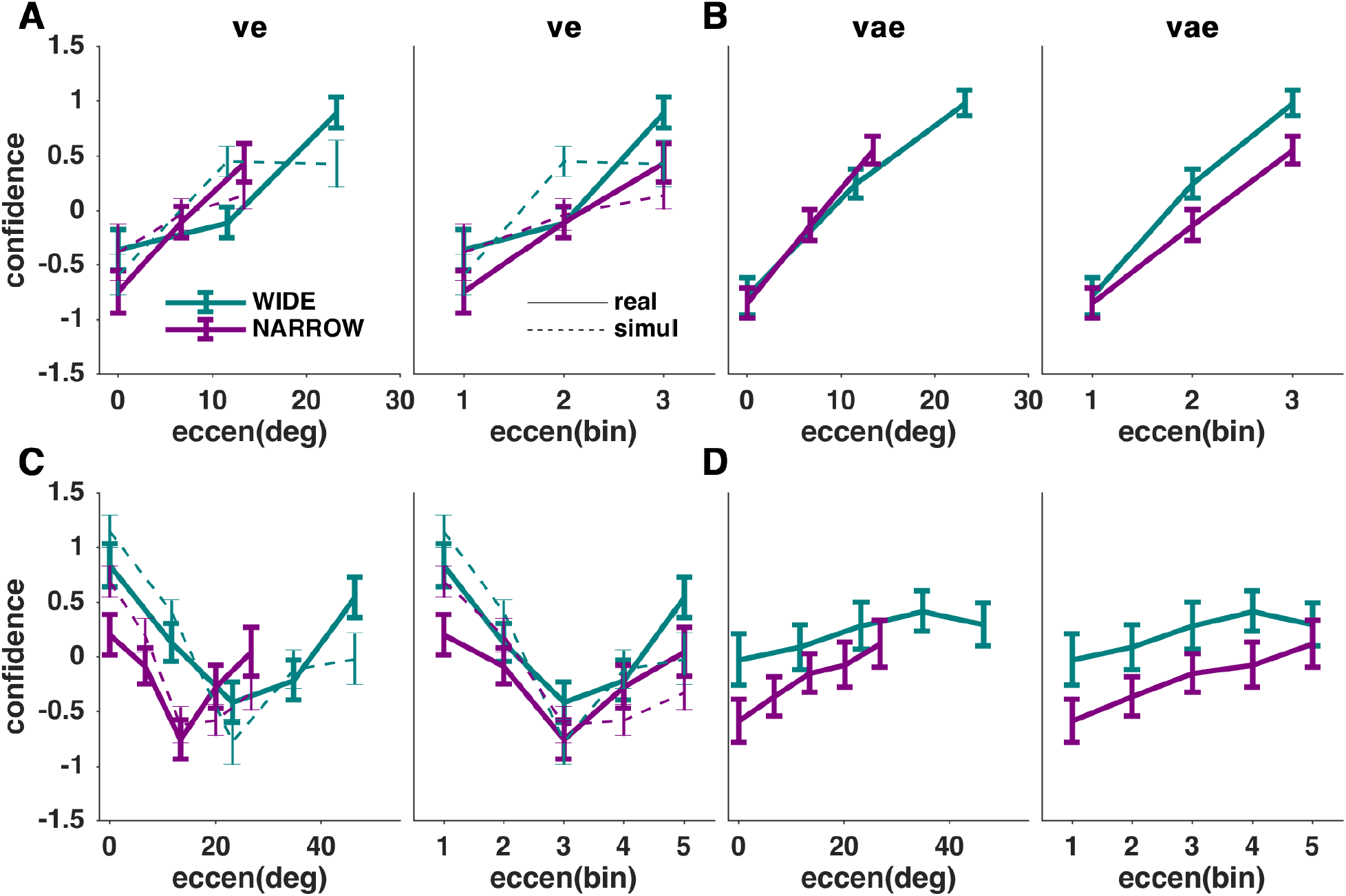
Confidence ratings of sound localization judgements. (A) Normalized confidence ratings in the AV trial for each context as a function of sound eccentricity in degrees (left) and normalized to the same range (right). The solid graph displays the group-mean and standard error of mean for the actual data (n=21), dashed lines the confidence ratings obtained from the Bayesian Causal inference model using model selection a decision function. (B) Confidence ratings for the A trial. (C) Confidence rating as a function of the audio-visual discrepancy, ΔVA, showing the actual data (solid) and the model prediction (dashed). (D) Same as (C) but for the A trial data. Teal: WIDE; Plum: NARROW.

For the AV trials, we expected confidence to also vary with the magnitude of the multisensory discrepancy, ΔVA. Such a dependency is directly predicted by the BCI models, as the posterior likelihood distribution of each position estimate becomes wider when the physical discrepancy increases (Figure 5C, dashed lines). Given that we had no specific expectation about a parametric scaling of confidence with ΔVA that would be consistent between ve and vae biases, we analysed this dependency using a repeated-measures ANOVA. For the AV trial, this provided decisive evidence for an effect of ΔVA (F_(4,200)_=5.9, p=10^−5^, BF=10^4^), anecdotal evidence for an effect of context (F=5.99, p=0.01, BF=1.2) and decisive evidence against an interaction (F=0.76, p=0.54, BF=−106).

For the A trial, we would expect confidence to vary with the previous ΔVA, if the hypothesis is correct that recalibration and integration biases are both driven by the same causal inference process. However, if the alternative hypothesis is correct that recalibration is driven by the belief that the unisensory modality is biased, one should expect confidence to scale with the contextual manipulation only; a larger context, hence a wider spacing between putative sound locations, should increase participants’ confidence about where the sound was originating from. The data provided no evidence for an effect of ΔVA (F=0.82, p=0.51, BF=−2.1), substantial evidence against an interaction between ΔVA and context (F=0.27, p=0.89, BF=−5.4) and very strong evidence for an effect of context (F=4.0, p=0.04, BF=60). Hence, for integration these confidence ratings are in agreement with Bayesian models of sensory causal inference, while for the aftereffect they point to an influence of spatial context but no influence of multisensory discrepancy.

### Comparison of aftereffect bias here and in previous work

One may wonder whether the data obtained in this experiment were somehow untypical and hence would not allow seeing a change in aftereffect with context. We hence directly compared the magnitude of the aftereffect biases obtained here to data obtained in similar versions of this task in our previous work. To this end we reanalysed the data from a previous EEG study (Park & Kayser, 2021) (n=19), a study manipulating time delays in between trials (n=41) or introducing a mask between AV and A trials (n=44)(Park & Kayser, 2020) and a dataset obtained in a study comparing young and older participants (n=22; younger participants only) (Park et al., 2021). These paradigms differed in their number (using 9, 5, 9 and 9 levels respectively) or range of spatial discrepancies used (34°, 24°, 32° and 32°). For each of these and the current data (n=42 across contexts) we obtained an estimate of the aftereffect magnitude as the average vae bias computed over the largest two levels of discrepancies per experiment. The resulting study-averaged aftereffect magnitudes were comparable (mean±s.e.m.: 1.2±0.2, 0.83±0.12, 1.23±0.17, 2.1±0.37 for the previous studies and 0.97±0.11 for the present data) and a two-sample Kolmogorov-Smirnov test did not provide evidence that the distributions of aftereffects differed (K=0.16, p=0.31).

## Discussion

By manipulating the spatial range over which audio-visual stimuli were presented in a ventriloquism paradigm we found that sensory integration (the ve bias) but not recalibration (the immediate aftereffect bias, vae) changed with the experienced context of multisensory discrepancies. When stimuli were paired over a larger range of spatial discrepancies the integration bias became stronger, and the combined behavioural and modelling evidence suggests that this enhanced integration bias arises from an enhanced a priori binding tendency in the wider context. This shows that multisensory integration adapts to the momentary sensory context, while the immediate recalibration bias seems to be largely independent of this.

### The integration bias adjusts based on the experienced causal relations

Multisensory integration is guided by an implicit judgement of the likely causal relation between two sensory cues, leading to a stronger binding when two cues are assumed to originate from a common source (Odegaard et al., 2017; Parise et al., 2012; Rohe & Noppeney, 2015b; Wallace et al., 2004). In a typical ventriloquism paradigm, the two sensory cues are presented in a pseudo-randomized fashion and the observer estimates a presumed degree of general causal relation from the overall pattern of spatio-temporal regularity in their co-occurrence. We reasoned that the a priori binding tendency underlying such causal judgements should adapt to a given spatial context, for example based on the range of observed discrepancies. Especially, for audio-visual stimuli with the same relative reliabilities we expected that pairing these over a larger range of the horizontal plane should facilitate the belief of a common underlying cause, because the signal to noise ratio for the localization of each pair (the ratio of sensory reliability to the range of spatial discrepancies) would be enhanced. To probe this, we presented the same audio-visual stimulus pairs from the same number of discrete locations and using the same timing in two contexts that differed only in the range of experienced multisensory discrepancies. Our data show that indeed the ventriloquism bias became stronger with larger spatial context. The Bayesian causal inference model directly supported these conclusions, as the behavioural data was better explained by a model that adapts to the contextual manipulation and does so by assigning a higher a priori binding tendency to the wider context.

Previous work as shown that the a priori binding tendency is malleable, varies within an individual between tasks (Odegaard & Shams, 2016) and can be altered by brief exposure to spatio-temporally incongruent stimuli (Odegaard et al., 2017), while it is rather stable to changes in selective attention (Odegaard et al., 2016). All in all, these studies show that the degree to which multisensory signals influence each other during perceptual judgments depends not only on their individual reliability and their apparent discrepancy, but also on the overall spatio-temporal pattern of their co-occurrence (Noppeney, 2021; Parise et al., 2012; Wallace et al., 2004). The latter can be experimentally manipulated by exposure to distinct contexts such as done here, or by specific training paradigms (Habets et al., 2017; Tong et al., 2020). Together with recent work suggesting that the a priori binding tendency is also updated based on the sensory evidence in the immediate past (Rohe et al., 2019) this shows that the processes underlying multisensory binding are similarly adaptive as other perceptual processes typically studied in unisensory contexts.

### No evidence for the aftereffect bias to adjust to context

In contrast to the integration bias, the aftereffect bias did not differ between contexts and the statistical analyses provided decisive evidence for a null result. This finding has interesting implications for the discussion of computational underpinnings of the aftereffect. Previous experiments have shown that the aftereffect can emerge independently on an immediate (trial-by-trial) scale and over cumulative long-term exposure (Bruns & Röder, 2015, 2019; Park & Kayser, 2021; Wozny & Shams, 2011b). Previous work has speculated that the immediate aftereffect may be directly related to the integration in the preceding trial, whereby the (implicit) judgement about a putative causal relation between two sensory cues moulds the recalibration process (Badde et al., 2020; Bruns & Röder, 2015; Noppeney, 2021; Park & Kayser, 2019; Rohlf et al., 2021). Indeed, studies have shown that the recalibration bias is directly related to the direction and the degree of the previous multisensory discrepancy rather than e.g. to the relative location difference between the unisensory test stimulus and a previous unisensory stimulus (Park & Kayser, 2019; Wozny & Shams, 2011b). Furthermore, recalibration is facilitated by distributed attention, possibly by strengthening the a priori binding tendency and thereby facilitating both integration bias and the (cumulative) aftereffect (Badde et al., 2020). In our previous work, a statistical relation between the aftereffect and the integration biases was observed consistently across multiple datasets, and in the present data we confirmed that the integration bias in the preceding multisensory trial adds predictive power for the aftereffect bias over and on top of the sensory discrepancy. However, neither the statistical influence of discrepancy nor that of the previous integration bias differed between contexts. Importantly, we ensured that the obtained aftereffect biases in the present study were not unusually small and hence may obscure any context effect. This speaks against a strict link between multisensory integration in the AV trial and the immediate aftereffect in the subsequent A trial.

An alternative viewpoint holds that the recalibration bias is independent of whether the preceding multisensory cues were deemed causally related, but rather serves to adjust behaviour dependent on the belief of a modality-specific bias (Di Luca et al., 2009; Noppeney, 2021; Zaidel et al., 2013). This belief in a modality-specific bias may still be shaped by the patterns of spatio-temporal regularity in the sensory environment and the experienced unisensory reliabilities, given that obtaining a direct estimate of the accuracy of each individual modality’s representation requires external feedback. If this account of the aftereffect bias is correct, one would not expect a difference in aftereffect bias between contexts. This is precisely what we observed in the present data.

Based on a similar paradigm we have previously reported evidence for a common neural substrate for the integration bias and the immediate aftereffect in neuroimaging (MEG) data (Park & Kayser, 2019, 2021). Specifically, these previous data support the hypothesis that medial parietal regions may contribute to the immediate ventriloquism aftereffect based on the maintenance of fused multisensory representations between trials. The present data do not rule out that a shared neural substrate underpins both integration and recalibration, as both were statistically related beyond the immediate influence of the multisensory discrepancy. However, the present data suggest that any presumed role of parietal regions in recalibration may not be the sole determinant of the immediate aftereffect bias, a conclusion also supported by a previous study revealing no effect of mnemonic manipulations on the immediate aftereffect (Park & Kayser, 2020).

### Confidence ratings suggest a high-level contribution to the aftereffect bias

As previous work offered no clear prediction as to whether the immediate aftereffect would change with the contextual manipulation, we also obtained subjective confidence ratings for the localization judgements. Confidence has proven to be a useful tool to dissociate sensory from decision-level origins of serial or adaptive effects in unisensory perceptual tasks (Benwell et al., 2019; Cicchini et al., 2018; Fleming & Lau, 2014; Gallagher et al., 2019). In particular, any dissociation of how response biases and confidence scale with task manipulations can provide insights about their computational origin in the sensory-decision process.

For the ventriloquism bias we found that confidence scaled with task difficulty and with the contextual manipulation, as predicted by the Bayesian causal inference model. For the aftereffect trials confidence did not vary with the degree of multisensory discrepancy, but varied with context. We take this as further evidence that the aftereffect bias is not only a sensory-level effect, as this would imply a similar scaling of aftereffect bias and confidence with discrepancy (Gallagher et al., 2019). Rather, the observation that confidence, but not bias, changes with context points to contributions from higher level and post-sensory processes in the aftereffect.

## Conclusion

Our results show that multisensory binding is similarly adaptive as many aspects of unisensory perception. The overall belief that two sensory cues are causally related is shaped by the context-specific spatio-temporal patterns of stimulus co-occurrence. This context-specific causal inference directly results in an adaptive integration bias by which one stimulus is misjudged in relation to the other in a context-sensitive manner. In our data this change in the belief of a causal relation did not translate into an adapted ventriloquism aftereffect, suggesting that this bias of unisensory perceptual judgements following a multisensory stimulus is not a simple or a pure sensory-level consequence of the multisensory integration process.

## Acknowledgements

This work was supported by the European Research Council (to C.K. ERC-2014-CoG; grant No 646657). We would like to thank a number of research assistants for their help during data collection.

## AUTHOR CONTRIBUTIONS

CK and HP Conceived study; collected and analyzed data; wrote the manuscript.

## ADDITIONAL INFORMATION

Authors declare no competing interests.

